# A lack of repeatability creates the illusion of a trade-off between basal and plastic cold tolerance

**DOI:** 10.1101/2021.09.24.461715

**Authors:** Erica O’Neill, Hannah E. Davis, Heath A. MacMillan

## Abstract

The thermotolerance-plasticity trade-off hypothesis predicts that ectotherms with greater basal thermal tolerance have a lower acclimation capacity. This hypothesis has been tested at both high and low temperatures but the results often conflict. If basal tolerance constrains plasticity (e.g. through shared mechanisms that create physiological constraints), it should be evident at the level of the individual, provided the trait measured is repeatable. Here, we used chill-coma onset temperature and chill-coma recovery time (CCO and CCRT; non-lethal thermal limits) to quantify cold tolerance of *Drosophila melanogaster* across two trials (pre- and post-acclimation). Cold acclimation improved cold tolerance, as expected, but individual measurements of CCO and CCRT in non-acclimated flies were not (or only slightly) repeatable. Surprisingly, however, there was still a strong correlation between basal tolerance and plasticity in cold-acclimated flies. We argue that this relationship is a statistical artefact (specifically, a manifestation of regression to the mean; RTM) and does not reflect a true trade-off or physiological constraint. Thermal tolerance trade-off patterns in previous studies that used similar methodology are thus likely to be impacted by RTM. Moving forward, controlling and/or correcting for RTM effects is critical to determining whether such a trade-off or physiological constraint truly exists.

## Introduction

Thermotolerance is widely considered a major determinant of ectotherm geographic distribution, as thermal limits— particularly lower thermal limits— can strongly predict ectotherm range limits [1–3]. In an era of global climate change [4], the ongoing persistence of ectotherms already residing near their thermal limits will depend on the ability of these organisms to adapt or respond plastically to changing thermal conditions [5,6]. Consequently, understanding evolutionary and physiological constraints on ectotherm cold tolerance adaptation and plasticity may be vital for predicting how ectotherms will be affected by global climate change [5].

Cold tolerance is the ability of an organism to survive exposure to low temperature extremes [7]. In many ectotherms, exposure to sublethal low temperatures physiologically primes the organism to better respond to subsequent cold stresses (reviewed for insects in: [8,9]). Consequently, cold tolerance can be conceptually divided into two categories: basal tolerance (an organism’s baseline level of cold tolerance; e.g. [10,11]), and induced tolerance (an organism’s level of cold tolerance following some adjustment period; e.g. [10]). Physiological priming that results in an improvement in thermotolerance is a form of phenotypic plasticity and is typically named according to the length of the sublethal temperature exposure, from short-term (minutes-hours) “hardening” to long-term (days-weeks) “acclimation” [12,13].

Basal and plastic cold tolerance can be quantified using a variety of metrics in chill-susceptible insects (reviewed in [8]). For instance, critical thermal minimum (CT_min_) is the temperature, upon cold exposure, at which neuromuscular coordination of the organism becomes impaired [14]. If cold exposure persists, CT_min_ is followed by chill-coma onset (CCO), the temperature at which neuromuscular function is entirely lost, resulting in complete paralysis [14]. Following CCO and a period of time in the cold, chill-coma recovery time (CCRT) is the time taken for an organism to regain neuromuscular function after being removed from coma-inducing conditions [15]. For all three metrics— CT_min_, CCO, and CCRT— low values indicate high cold tolerance. In contrast to the above non-lethal metrics, lethal measures of cold tolerance include survivorship (the proportion of organisms surviving some cold exposure; e.g. [11]), lower lethal temperature (LLT; the temperature at which a certain proportion of individuals die under a fixed duration of cold exposure (this is calculated using survivorship); e.g. [16]), and lower thermal limit (LTL; an average temperature of mortality; e.g. [17]).

One critical uncertainty and point of contention among thermal biologists is whether basal thermotolerance constrains plastic thermotolerance, such that animals with greater basal tolerance— either cold tolerance (as described above) or heat tolerance— have a lower capacity for acclimation [11,16–22]. To address this question, several researchers have used thermotolerance metrics to test for a trade-off between basal and induced cold tolerance–called a tolerance-plasticity trade-off (summarized in Table 1). While some studies (spanning a variety of taxa) have documented evidence for such a trade-off at low temperatures (e.g. [11,20,22]), others have found partial or no such evidence [16,17,21]. Similarly, in a recent review of heat tolerance-plasticity trade-off studies, approximately half of the studies gathered— 17 out of 30— did not support a trade-off [23].

**Table 1.**
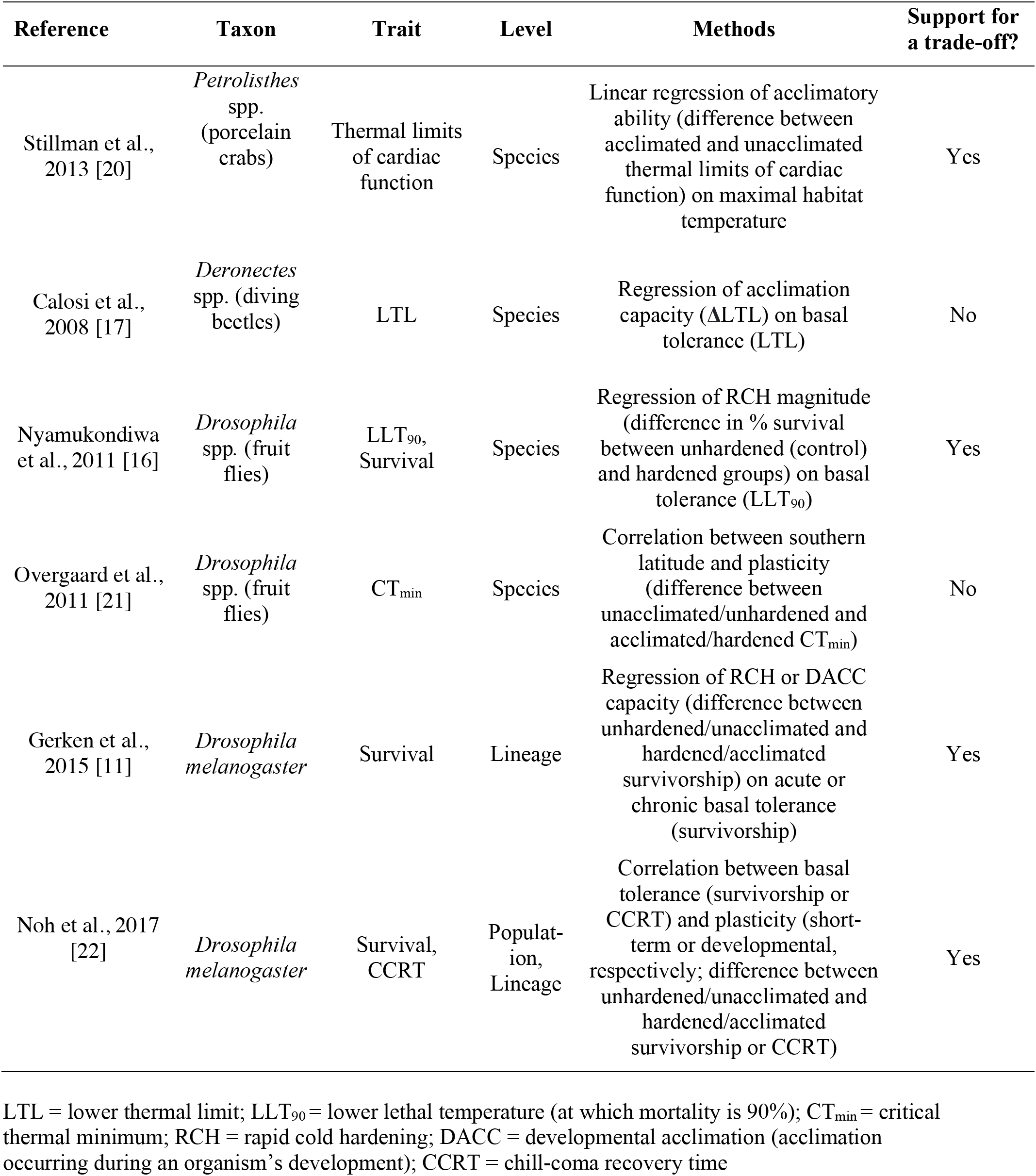
Examples of studies that have tested for a trade-off between basal cold tolerance and plasticity. “Support for a trade-off” refers only to results for a trade-off tested at low temperatures (some of the following studies also test for a trade-off at high temperatures).

| Reference | Taxon | Trait | Level | Methods | Support for a trade-off? |
| --- | --- | --- | --- | --- | --- |
| Stillman et al., 2013 [20] | *Petrolisthes* spp. (porcelain crabs) | Thermal limits of cardiac function | Species | Linear regression of acclimatory ability (difference between acclimated and unacclimated thermal limits of cardiac function) on maximal habitat temperature | Yes |
| Calosi et al., 2008 [17] | *Deronectes* spp. (diving beetles) | LTL | Species | Regression of acclimation capacity (ΔLTL) on basal tolerance (LTL) | No |
| Nyamukondiwa et al., 2011 [16] | *Drosophila* spp. (fruit flies) | LLT<sub>90</sub>, Survival | Species | Regression of RCH magnitude (difference in % survival between unhardened (control) and hardened groups) on basal tolerance (LLT<sub>90</sub>) | Yes |
| Overgaard et al., 2011 [21] | *Drosophila* spp. (fruit flies) | CT<sub>min</sub> | Species | Correlation between southern latitude and plasticity (difference between unacclimated/unhardened and acclimated/hardened CT<sub>min</sub>) | No |
| Gerken et al., 2015 [11] | *Drosophila melanogaster* | Survival | Lineage | Regression of RCH or DACC capacity (difference between unhardened/unacclimated and hardened/acclimated survivorship) on acute or chronic basal tolerance (survivorship) | Yes |
| Noh et al., 2017 [22] | *Drosophila melanogaster* | Survival, CCRT | Population, Lineage | Correlation between basal tolerance (survivorship or CCRT) and plasticity (short-term or developmental, respectively; difference between unhardened/unacclimated and hardened/acclimated survivorship or CCRT) | Yes |

LTL = lower thermal limit; LLT<sub>90</sub> = lower lethal temperature (at which mortality is 90%); CT<sub>min</sub> = critical thermal minimum; RCH = rapid cold hardening; DACC = developmental acclimation (acclimation occurring during an organism’s development); CCRT = chill-coma recovery time

Importantly, the studies described above and in Table 1 tested for a trade-off between some measure of basal cold tolerance and plasticity predominantly at the species, lineage, or population level. But if basal tolerance constrains plasticity through shared physiological mechanisms of tolerance, such a pattern should arise at the level of the individual. The CT_min_, for example, is thought to occur as a result of sudden ionoregulatory failure (spreading depolarization) in the nervous system [24,25], so, for example, constitutive expression of ionoregulatory proteins (e.g. ion channels) that help to lower the basal CT_min_ may approach the “ceiling” to expression that can occur during thermal acclimation. Basal and induced cold tolerance have similarly been suggested to share renal mechanisms that help prevent ionoregulatory collapse (at least in *Drosophila;* [26–28]). We hypothesized that if such constraints exist, they should be evident at the individual level and may underlie trade-off patterns previously observed at higher levels of organization. To our knowledge, however, only three studies have ever investigated the relationship between basal thermotolerance and plasticity at the individual level in ectotherms, each time at high temperatures [29–31]. Thus, turning our attention away from species/lineage/population level comparisons and towards individual variation— which has historically been underutilized in physiology [32,33]— may prove informative.

Here, we tested whether basal cold tolerance constrains plasticity at the individual level in *Drosophila melanogaster*. We did so by quantifying cold tolerance in individual *D. melanogaster* across two trials (pre- and post-cold-acclimation; Fig. 1). Pre-acclimation measurements give an estimate of basal cold tolerance, while post-acclimation measurements give an estimate of induced cold tolerance. If basal cold tolerance constrains plasticity, we expected that individuals with higher basal cold tolerance (lower trial 1 measurements) would tend to have diminished acclimation capacity (a reduced difference in thermotolerance between trial 1 and trial 2) relative to individuals with lower basal cold tolerance. Such methodology, where acclimation capacity is correlated with basal tolerance and is measured as the difference between basal and induced tolerance, is a common method of testing the tolerance-plasticity trade-off hypothesis (e.g. [11,16,17,22]; Table 1). Because different cold tolerance traits have different underlying mechanisms and vary in different ways among and within populations and lineages of *Drosophila* [3,8,34,35], we carried out our experiment twice using different metrics of thermotolerance: chill-coma onset (CCO) and chill-coma recovery time (CCRT; Fig. 1).

**Figure 1.**
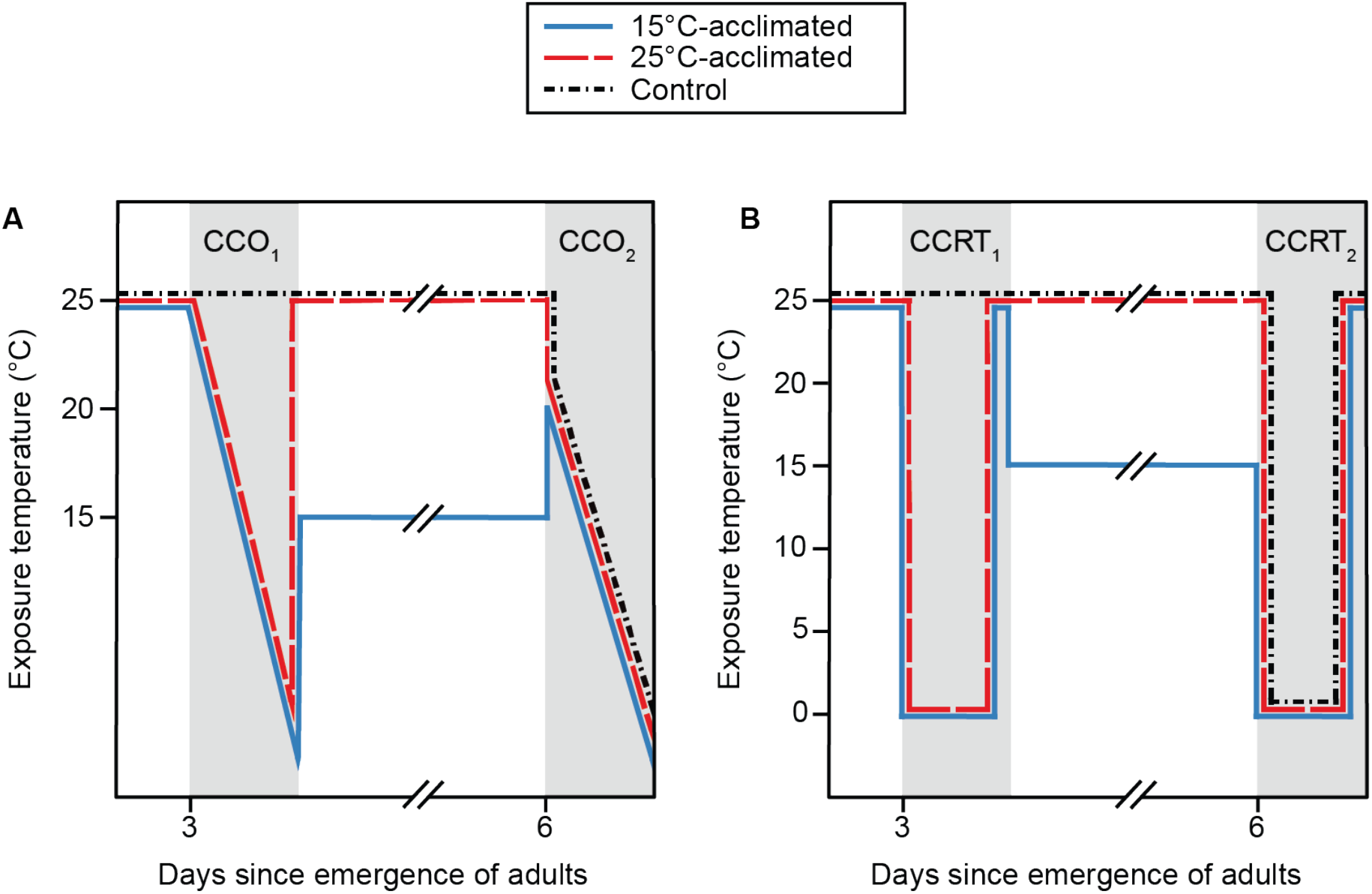
Outline of experimental design for CCO (A) and CCRT (B). All flies (female *D. melanogaster*) were reared at 25°C. The 15°C- and 25°C-acclimated groups underwent an initial thermotolerance measurement (CCO_1_ or CCRT_1_; pre-acclimation), and were subsequently kept at 15°C or 25°C, respectively, for the next three days; the control group was kept at 25°C throughout this time. All three groups underwent the second thermotolerance measurement (CCO_2_ or CCRT_2_; post-acclimation).

## Materials and Methods

### Experimental overview

Chill-coma onset and chill-coma recovery time (CCO and CCRT; non-lethal metrics of thermotolerance) were used to quantify cold tolerance of individual flies across two trials (pre- and post-acclimation; Fig. 1) to determine the relationship between basal tolerance and acclimation capacity in cold-acclimated flies (“15°C-acclimated”; Fig. 1). Two other groups of flies were included in the experiment: 1) flies that were not subject to cold acclimation between trials (“25°C-acclimated”); 2) flies that were not subject to cold acclimation and that underwent trial 2 but not trial 1 (“Control”; Fig. 1). The 25°C-acclimated flies were used to assess the within-individual repeatability (hereafter, “repeatability”) of CCO and CCRT in the absence of cold acclimation, thereby estimating the repeatability of basal tolerance as measured by both metrics. Repeatability is a descriptor of the within-individual consistency, or predictability, of a trait over repeat measurements [45,46]. Repeatability of CCO and CCRT in non-cold-acclimated flies is essential to addressing the trade-off hypothesis because otherwise measurements of CCO and CCRT are unreliable indices of basal thermotolerance. Unhelpfully, the repeatability of ectotherm thermotolerance traits is infrequently studied— repeatability has in fact never been estimated for CCO and has only been estimated twice for CCRT (with conflicting results, and neither time in *Drosophila*; [36,37]) — providing little hint of what to expect. Though, of the ectotherm thermotolerance traits studied, most show at least moderate repeatability (Table 2). Importantly, if trial 1 had some effect on trial 2 measurements, estimates of the repeatability of CCO and CCRT in 25°C-acclimated flies (who by necessity underwent both trials) would not necessarily be accurate representations of the repeatability of basal tolerance. Thus, control flies (“Control”; Fig. 1) were included to check whether trial 1 itself influenced thermotolerance in trial 2.

**Table 2.** Summary of studies that have estimated within-individual repeatability of thermotolerance traits in ectotherms. “Repeatable?” refers to whether the repeatability of the trait in question is statistically significant (that is, significantly different from 0); repeatability ranges from 0 (no repeatability) to 1 (high repeatability). Adjusted repeatability controls for group-wide shifts in trait values from trial to trial while non-adjusted repeatability does not. Search methods for finding the gathered studies are outlined in the Table S1.

| Reference | Trait | Taxon/Taxa | Repeatable? | Repeatability | Method of estimating repeatability |
| --- | --- | --- | --- | --- | --- |
| Andrew and Kemp, 2016 [36] | CCRT | *Eurema smilax* (small grass yellow butterfly) | Yes | 0.4; 0.405 | Adjusted (Pearson correlation coefficient); non-adjusted (Intraclass correlation coefficient) |
| Scharf et al., 2016 [37] | CCRT | *Myrmeleon hyalinus* (antlion); *Vermileo* sp. (wormlion) | Yes; No | 0.306; 0.198 | Adjusted (Pearson correlation coefficient) |
| Hawes, 2006 [38] | SCP | *Cryptopygus antarcticus* (Antarctic springtail) | Yes | 0.910 | Adjusted (Spearman’s rank correlation) |
| Worland, 2005 [39] | SCP | *Tullbergia antarctica* (sub-Antarctic springtail) | Yes | 0.98 | Adjusted (methods unspecified) |
| Ditrich, 2018 [40] | SCP | *Pyrrhocoris apterus* (linden bug) | Yes | 0.61-0.84 | Adjusted (Pearson correlation coefficient) |
| Morgan et al., 2018 [29] | CT<sub>max</sub> | *Danio rerio* (zebrafish) | Yes | 0.447* | Adjusted (GLMM-based repeatability) |
| Bard and Kieffer, 2019 [41] | CT<sub>max</sub> | *Acipenser brevirostrum* (shortnose sturgeon) | Yes | N/A | Linear regression between first and second measurements |
| O’Donnell et al., 2020 [42] | CT<sub>max</sub> | *Salvelinus fontinalis* (brook trout) | Yes | 0.48 | Adjusted (LMM-based repeatability) |
| Grinder et al., 2020 [43] | CT<sub>max</sub> | *Poecilia reticulata* (Trinidadian guppy) | Yes | 0.43 | Adjusted; non-adjusted (GLMM-based repeatability) |
| Claireaux et al., 2013 [44] | UILT | *Dicentrarchus labrax* (European sea bass) | Yes | 0.35-0.68 | Adjusted (Pearson correlation coefficient) |

\*when first trial was omitted. CCRT = chill-coma recovery time; SCP = supercooling point; CT<sub>max</sub> = critical thermal maximum; UILT = upper incipient lethal temperature; LMM = linear mixed-effects model; GLMM = generalized linear mixed-effects model

### Animal husbandry

*Drosophila melanogaster* used in this study originate from 35 isofemale lines originally captured in London and Niagara on the Lake, Ontario, Canada [47]. Flies were reared in a 25°C incubator (12 h: 12 h light:dark cycle) in 250 mL bottles containing ~50 mL of a banana, corn syrup, and yeast-based medium. To gather new eggs, adult flies were transferred to bottles containing fresh media (~100 per bottle) and allowed to lay eggs for approximately 2 h before being removed. The eggs laid in this time (~150 per bottle) were reared at 25°C until adult emergence (~10 days). On their day of emergence, adults were transferred to new bottles (such that all flies were within 25 h of age), where they remained for ~24 h at 25°C to ensure all females were mated before flies were anaesthetized (<15 min exposure to CO_2_) and sorted by sex. It was important to ensure females were mated because mating status can affect cold tolerance in insects (e.g.[48]).

Approximately 30 females (per run of the experiment) were then placed individually into 1.7 mL microcentrifuge tubes. Prior to this, the microcentrifuge tubes were prepared by puncturing one small hole in each tube lid and filling the tube with ~0.5 mL of the rearing diet. Early trials confirmed that the food inside these tubes stayed well hydrated for at least 72 h and thus would not cause desiccation stress. The microcentrifuge tubes were placed in a 25°C incubator where they remained for typically 48-50 h (in one CCO run, flies remained for 53 h) before the first thermotolerance measurement (trial 1). This delay was to prevent CO_2_ exposure from interfering with cold tolerance [49]. After trial 1 (pre-acclimation), flies were placed into new microcentrifuge tubes (as described above) and held for three days at either 15°C or 25°C until trial 2 (post-acclimation). Control flies did not undergo trial 1 measurements but were moved into fresh microcentrifuge tubes while, or directly after, the trial was carried out (Fig. 1).

### Measurement of chill-coma onset

Chill-coma onset (CCO) was measured by placing flies individually into 3.7 mL screw-top glass vials, attaching these vials to an aluminum rack, and submerging the rack in a custom built cooling bath (1:1 v/v water:ethylene glycol; as in [50]). The temperature of the cooling bath began at 25°C (for trial 1) or 20°C (for trial 2) and decreased at a rate of 0.1°C/min. The temperature was monitored by three type-K thermocouples placed at different locations in the bath. After the temperature of the bath reached ~10°C, the glass vials were tapped frequently with a metal rod to stimulate movement in the flies [27]. CCO was recorded as the temperature at which a fly stopped moving and was unresponsive to tapping on the vial. The temperature decreased until all flies had entered chill-coma, at which point the rack with vials was removed from the bath. During trial 1, control flies were transferred into 3.7 mL screw-top glass vials and kept in a 25°C incubator for the duration of the trial (~4 h) before being transferred into fresh microcentrifuge tubes (to maintain similar levels of handling for the control and non-control flies).

### Measurement of chill-coma recovery time

Chill-coma recovery time (CCRT) was measured by placing flies individually into 3.7 mL screw-top glass vials and placing these vials in a water-ice mixture within a Styrofoam box inside a fridge (such that flies were exposed to ~0°C). The vials remained in this setup for 6 h, after which they were removed from the water-ice mixture and set on a counter at room temperature to record CCRT. CCRT was recorded as the time taken– from the moment the vials were removed from the water-ice mixture– for flies to stand upright on all six limbs [27]. To control for handling, control flies were transferred into fresh microcentrifuge tubes while trial 1 was being carried out. We also recorded room temperature during our CCRT recordings. Room temperature varied among experimental runs (mean ± sd: 24.5 ± 1.3°C), and one run was discarded and then replaced because CCRT tended to be slower (although not significantly) and room temperature on that day was lower (21.9°C) than all other runs. Ultimately, room temperature had no significant effect on CCRT, and was shared among all flies in each run (and we included flies from all three treatment groups in each run), so we excluded it as a factor in downstream analyses.

### Data analysis

All data analyses were carried out in R version 3.6.2 [51]. Individuals that were lost or accidentally crushed between trials were removed from the analysis. To test whether treatment (15°C vs. 25°C acclimation) affected CCO or CCRT (that is, induced plasticity), linear mixed-effects models were performed (fixed effects: trial (1 vs. 2), treatment group; random effects: run date, individual ID (nested within run date)). Repeatability of CCO or CCRT in 25°C-acclimated flies was estimated using the rpt() function in the R package rptR (unadjusted repeatability with a normal distribution of data assumed; number of parametric bootstraps = 1000 [52]). To test whether disturbances to the flies associated with carrying out trial 1 affected trial 2 measurements, trial 2 measurements (CCO_2_ or CCRT_2_) of the control and 25°C-acclimated groups were compared using linear mixed-effects models (fixed effect: treatment group; random effect: run date). Finally, to assess the relationship between basal tolerance (CCO_1_ or CCRT_1_) and acclimation capacity (CCO_1_ – CCO_2_ or CCRT_1_ – CCRT_2_), two more linear mixed-effects models were carried out (fixed effects: CCO_1_ or CCRT_1_, treatment group; random effect: run date); slopes of the 15°C- and 25°C-acclimated groups were compared by checking whether there was a significant interaction between the effects of CCO_1_ or CCRT_1_ and treatment group (15°C- vs. 25°C-acclimated) on acclimation capacity. All linear mixed-effects models were carried out using the lme() function in the R package nlme [53].

## Results and Discussion

### 1. Cold acclimation induced plasticity

As anticipated, 15°C-acclimation induced plasticity in CCO and CCRT. There was a significant effect of trial (LMM: F_1,126_ = 97.9, p < 0.0001) and treatment (15°C- vs. 25°C-acclimation; LMM: F_1,120_ = 107.0, p < 0.0001) on CCO, and a significant interaction between these effects (LMM: F_1,126_ = 106.4, p < 0.0001; Fig. 2A). Likewise, there was a significant effect of trial (LMM: F_1,191_ = 18.0, p < 0.0001) and treatment (LMM: F_1,185_ = 34.0, p < 0.0001) on CCRT, and a significant interaction between these effects (LMM: F_1,191_ = 30.7, p < 0.0001; Fig. 2B). This improvement in cold tolerance with cold acclimation is consistent with current literature [8]. Having verified that 15°C-acclimation did indeed induce plasticity in this study system, we moved on to characterizing the relationship between basal tolerance and plasticity.

**Figure 2.**
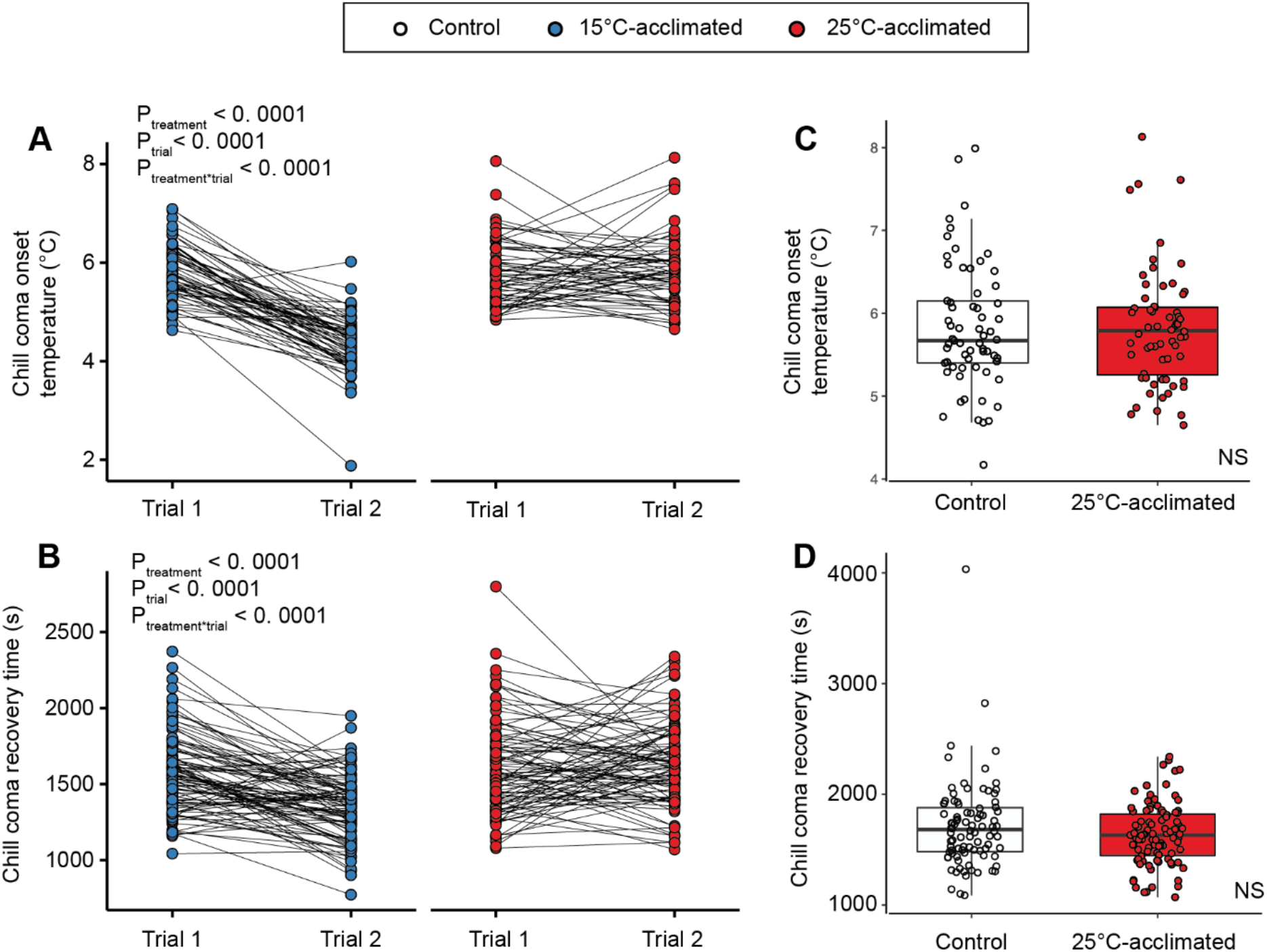
15°C-acclimation induced plasticity in CCO (A) and CCRT (B) in female *D. melanogaster*, and conditions in trial 1 did not affect subsequent measurements of CCO (C) or CCRT (D). Solid black lines in panels A and B connect an individual fly’s trial 1 measurement (CCO_1_ or CCRT_1_) to the same individual’s trial 2 measurement (CCO_2_ or CCRT_2_). The main effects of treatment and trial on CCO or CCRT were significant (p < 0.0001); there was a significant interaction effect between the two predictors (p < 0.0001). There was not a significant effect of treatment— control (did not undergo trial 1) vs. 25°C acclimation (did undergo trial 1)— on trial 2 measurements (CCO_2_ (C) or CCRT_2_ (D)).

### 2. Repeatability of CCO and CCRT in 25°C-acclimated flies was low

Earlier, within-individual repeatability was loosely defined as the consistency or predictability of a trait within individuals over repeated measurements [45,46]. More formally, repeatability (typically represented as “R”) is the proportion of total variance attributable to differences among (rather than within) subjects over repeat measurements of a trait [45,46]. “Subjects” may be any unit of measurement, from groups to— in this case— individuals. The non-repeatable portion of the total variance (1–R) is composed of measurement error and fluctuations of the trait within individuals [46,54]. We assumed that the lower the repeatability of CCO and CCRT in the 25°C-acclimated flies, the lower the likelihood of detecting a trade-off, if present, in the 15°C-acclimated flies. This is because low repeatability indicates that measurements of CCO and CCRT are unreliable indices of basal tolerance. This unreliability and consequential decreased likelihood of detecting a trade-off has two-fold reasoning. First, low repeatability of CCO and CCRT in the 25°C-acclimated flies — whether due to measurement error or “real” within-individual fluctuations in CCO or CCRT— indicates that single estimates of basal tolerance are not necessarily representative of a fly’s “true” (mean) basal tolerance and therefore may not predict a fly’s acclimation capacity. Second, and perhaps more importantly, since repeatability indicates the proportion of total variance attributable to differences among individuals [45,46], low repeatability means that we lack meaningful variation in basal tolerance that is theoretically necessary to detect a relationship between basal tolerance and plasticity. For both metrics, repeatability was, in fact, low: CCO in the 25°C-acclimated flies was not significantly repeatable (R ± SE = 0.14 ± 0.11, 95% CI = [0, 0.37], p = 0.1), while repeatability of CCRT in the 25°C-acclimated flies was significant, but low (R ± SE = 0.23 ± 0.09, 95% CI = [0.033, 0.41], p = 0.01).

Low repeatability in the 25°C-acclimated group may result if performing trial 1 had some effect (damage or acclimation) on the thermotolerance of flies in trial 2. If this is the case, repeatability in the 25°C-acclimated group does not give an accurate estimate of the repeatability of basal tolerance, as acclimated or damaged tolerance in trial 2 would not be considered representative of basal tolerance. However, there was no significant effect of treatment group (Control vs. 25°C-acclimated) on CCO_2_ (LMM: F_1,121_ = 0.2, p = 0.7; Fig. 2C). Similarly, there was no significant effect of treatment group on CCRT_2_ (LMM: F_1,177_ = 2.9, p = 0.09; Fig. 2D). This suggests that conditions experienced by flies during trial 1 did not affect subsequent measurements of thermotolerance. Thus, trial 2 measurements should be representative of basal tolerance (in the 25°C-acclimated flies) and repeatability of CCO and CCRT in the 25°C-acclimated flies should be accurate estimates of the repeatability of basal tolerance.

To some degree, low repeatability of CCO and CCRT is very likely due at least in part to measurement error, but because we are familiar with these techniques and the amount of measurement error that can be expected, we argue that true biological fluctuations and/or natural stochasticity in CCO or CCRT within individuals are primary contributing factors to the low repeatability we observed. Insect chill-coma has historically been associated with a reduction of neuromuscular excitability and depolarization of neuromuscular membrane potential [8]. More recent evidence has clarified the differing roles of the nervous and muscular systems in chill coma entry and recovery: entry into chill coma (CCO) appears to be caused primarily by depolarization of the central nervous system (CNS), while recovery from chill coma (CCRT) after hours in the cold involves rapid recovery of CNS function, followed by slower repolarization of muscle membrane potential as systemic ion balance is restored [55–57]. Within-individual stochasticity in mechanisms hypothesized to govern the maintenance of neuromuscular excitability and membrane potential at low temperatures— e.g. modulation of neuromuscular membrane composition (e.g.[58,59]), modulation of the thermal sensitivity of voltage-gated ion channels responsible for action potentials (e.g. [60,61]), or changes in ion transporters in neuromuscular cells [55,62] may contribute to low repeatability. CCRT is additionally influenced by an individual’s motivation to stand [3], so within-individual variability in motivation may lower repeatability of CCRT as well.

### 3. The illusion of a trade-off between basal and plastic cold tolerance

Given that estimates of the repeatability of basal tolerance were low for both CCO and CCRT, we did not expect to find a significant relationship between basal tolerance and acclimation capacity in the 15°C-acclimated flies. Surprisingly, however, we found a strong significant relationship between both CCO_1_ and acclimation capacity (CCO_1_ – CCO_2_) (F_1,118_ = 106.0, p < 0.0001) and CCRT_1_ and acclimation capacity (CCRT_1_ – CCRT_2_) (F_1,183_ = 239.4, p < 0.0001).

A more thorough consideration of the relationship between trial 1 and trial 2 measurements (particularly in the 25°C-acclimated flies) reveals how this significant trade off pattern arises in our data despite low— and in the case of CCO, no— repeatability of basal tolerance. In the 25°C-acclimated flies, there was a striking— and unexpected— relationship between trial 1 and trial 2 measurements within individuals: individuals that were more thermotolerant (lower CCO or CCRT) in trial 1 tended to be less tolerant in trial 2, and individuals that were less tolerant in trial 1 tended to be more tolerant in trial 2 (Fig. S1). One potential explanation for this pattern of data is that the least thermotolerant flies in trial 1 were cold-hardened by trial 1 conditions, while the most thermotolerant flies were ultimately damaged by trial 1 conditions. It is not illogical to imagine that an individual’s thermotolerance could subsequently improve because of exposure to low temperatures during trial 1. After all, hardening responses can be induced by only minutes-hours of sublethal cold exposure [13,63]. However, why the most tolerant flies in trial 1 would not only fail to cold-harden but ultimately be damaged by trial 1 conditions when seemingly less tolerant flies cold-hardened under the same conditions is unclear. Moreover, and more importantly, there is a better explanation for the pattern observed.

When repeated measurements are taken from the same subject, these measurements are not likely to be identical: some degree of random error— due to measurement error or fluctuation of the given trait— is almost inevitable [64]. Because of this random error, such repeated measurements are susceptible to a statistical phenomenon known as regression to the mean (RTM; [64,65]). Put simply, subjects that have more-extreme measurements in one trial will tend to have less-extreme measurements— measurements that are closer to their true mean— in the next trial. The pattern of within-individual variation we are attempting to explain in the 25°C-acclimated flies aligns with that expected from RTM (Fig. 4, S2). Due to this, and due to the near-inevitability of RTM when taking repeated measurements, we attribute the relationship between trial 1 and trial 2 measurements in the 25°C-acclimated flies to the effects of RTM, and not to some biological phenomenon. Rank plots (Fig. S2) further emphasize this unexpected positive correlation between basal thermotolerance and “acclimation capacity” in the 25°C-acclimated flies, as “acclimation capacity” (CCO_1_ – CCO_2_ or CCRT_1_–CCRT_2_) tends to be negative in flies with lower trial 1 measurements and positive in flies with higher trial 1 measurements, indicative of RTM.

Falsely ascribing significance (e.g. biological, economic, psychological) to manifestations of RTM is a common fallacy [65,66]. To avoid this error, one must separate effects expected from RTM, and effects (if any) beyond those expected from RTM [64]. In the 25°C-acclimated flies, differentiating such effects was straight-forward: there is no intervention applied or relevant biological phenomena expected to be occurring across the experimental period, so any apparent RTM effects can be reasonably attributed to RTM. In the 15°C-acclimated flies, on the other hand, both RTM and a thermotolerance-plasticity trade-off are expected to manifest as a positive correlation between basal tolerance and plasticity [64,65]. Thus, to determine whether the positive correlation previously noted in the 15°C-acclimated flies (Fig. 3) is truly evidence for a trade-off, we must determine how much of this correlation is attributable to RTM. In experimental studies, like ours, differentiation between the effects of RTM and other phenomena can be achieved quite easily with an adequate control group (i.e. a group for which repeat measurements are taken but no treatment/intervention is applied between trials) [65]. As RTM is expected to equally affect control and experimental groups, changes in the experimental group different from those observed in the control group suggest phenomena at play besides RTM. Happily, our 25°C-acclimated group acts as an ideal control for the 15°C-acclimated group in this regard.

**Figure 3.**
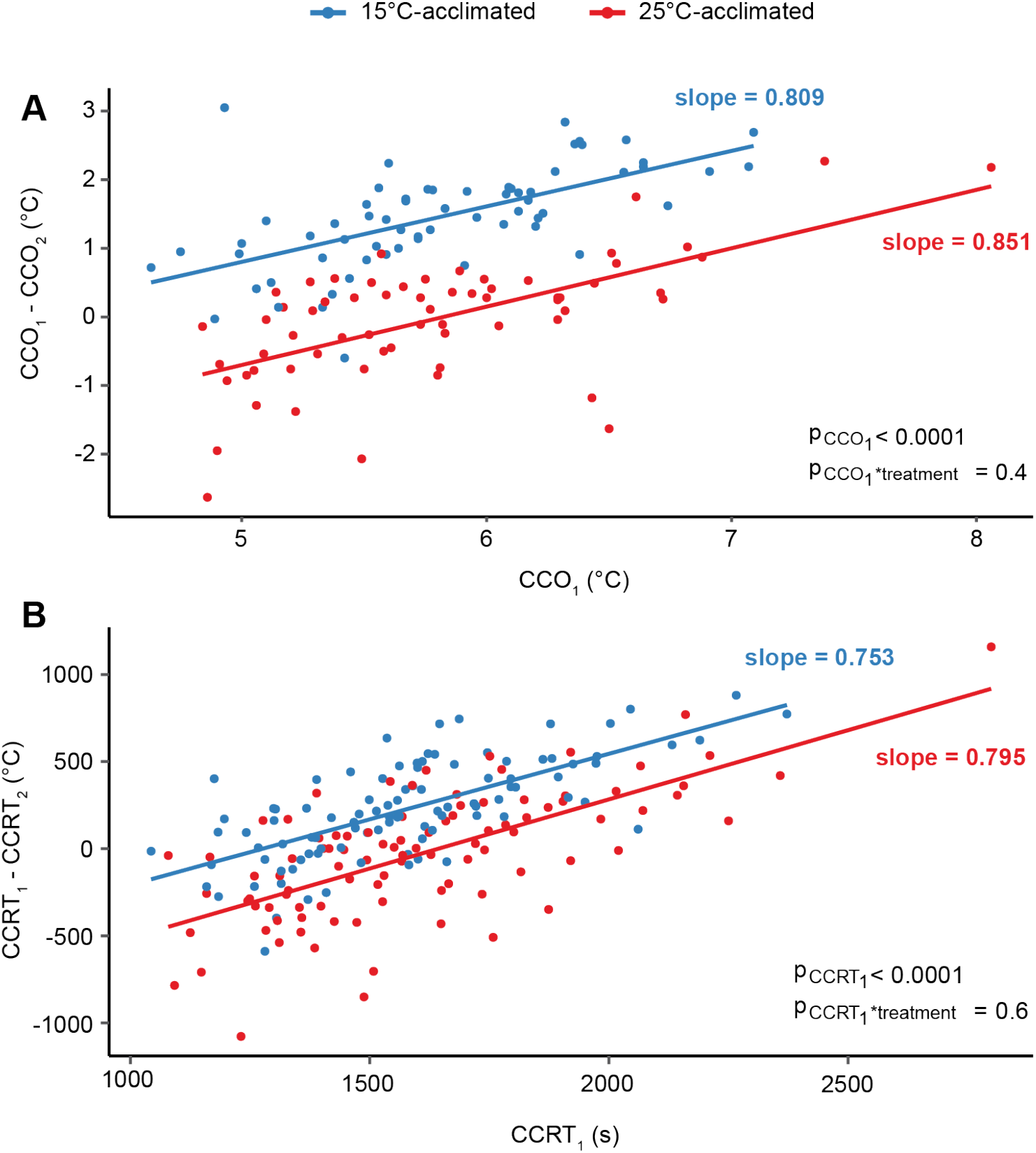
Relationship between CCO_1_ (A) or CCRT_1_ (B) and acclimation capacity (CCO_1_ – CCO_2_ or CCRT_1_–CCRT_2_) in female *D. melanogaster*. Fitted lines and slopes are based on a simple linear regression for each group for illustrative purposes (P-values derived from mixed effects models). A) CCO_1_ strongly predicted acclimation capacity in both groups (p < 0.0001). The slopes of the 15°C- and 25°C-acclimated groups were not significantly different (p = 0.4). B) CCRT_1_ strongly predicted acclimation capacity in both groups (p < 0.0001). The slopes of the 15°C- and 25°C-acclimated groups were not significantly different (p = 0.6).

### 4. The relationship between basal tolerance and acclimation capacity in the 15°C-acclimated flies is underlain by regression to the mean

If the slope between basal tolerance and “acclimation capacity” (CCO_1_ – CCO_2_ or CCRT_1_– CCRT_2_) in 25°C-acclimated flies is the same as the equivalent slope in the 15°C-acclimated flies, the apparent differential effect of cold acclimation on flies of high and low initial thermotolerance is not greater than the differential change expected from RTM. There was a significant relationship between CCO_1_ and acclimation capacity (CCO_1_ – CCO_2_) in both the 15°C- and 25°C-acclimated groups (F_1,118_ = 106.0, p < 0.0001; Fig. 3). The slopes of the relationships between CCO_1_ and acclimation capacity did not significantly differ between acclimation groups (F_1,118_ = 0.6, p = 0.4; Fig. 3A). Likewise, there was a significant relationship between CCRT_1_ and acclimation capacity (CCRT_1_–CCRT_2_) in both the 15°C- and 25°C- acclimated groups (F_1,183_ = 239.4, p < 0.0001), and the slopes of the relationships between CCRT_1_ and acclimation capacity similarly did not significantly differ between 15°C- and 25°C-acclimated flies (F = 0.2, p = 0.6; Fig. 3B). As the slopes of the 15°C- and 25°C-acclimated flies (Fig. 3) did not significantly differ for either thermotolerance metric, we cannot attribute the apparent differential effect of cold acclimation on flies of high or low initial thermotolerance to anything other than RTM. There is therefore no evidence for a tolerance-plasticity trade-off in the 15°C-acclimated flies.

### 5. How big of a problem is regression to the mean in studies of thermal tolerance?

When an independent variable (Y) contains the dependent variable (X) as a constituent (e.g. Y = X – Z; as when plasticity = basal tolerance – induced tolerance), these variables are not statistically independent [23,67]. Issues of interpretation resulting from this implicit statistical association have been articulated before (e.g. when estimating trade-offs between constitutive and induced defenses in plants [68]; when estimating costs of plasticity [67,69,70]; when assessing mass-dependent mass loss in birds [71,72]). Yet, this methodology has only recently been highlighted as a potential concern for thermotolerance-plasticity trade-off studies, and remains widespread [23]. Conclusions made on the basis of a relationship between baseline measurements and the difference between baseline and follow-up measurements may not necessarily be problematic if the null hypothesis for such an investigation is properly identified [71,72]. A proper null hypothesis should account for statistical bias resulting from phenomena like RTM [65]. However, there is very little consideration of RTM (or repeatability, for that matter) in previous thermotolerance-plasticity trade-off studies, precluding such identification (but see Deery et al. [30], who do not mention RTM but do address statistical bias).

It is unclear to what extent the results of past thermotolerance-plasticity trade-off studies are biased by RTM. We expect RTM to occur whenever repeated measurements of the same subjects are taken (as random error is unavoidable in experimental biology), whether the study is at the individual level or, as is the case for the vast majority of previous thermotolerance-plasticity trade-off studies, above the individual level [64,65] (but see: [29–31]). However, it is not clear under what specific methodological circumstances RTM occurs. For instance, if basal and induced tolerance are measured for the same group but different individuals within the group are used for each measurement, is RTM still expected to occur? This must be determined (e.g. through modeling hypothetical data) before effects of RTM can be corrected for (per Kelly and Price [65]).

In any case, the following certainly seems to be true: it is not yet known whether a thermotolerance-plasticity trade-off exists [23]. This lack of clarity stems from more than the inherent statistical association between basal tolerance and plasticity and issues of interpretation arising from it. For instance, many tolerance-plasticity trade-off studies are carried out on field organisms, meaning that environmental and genetic effects on thermotolerance cannot easily be unwoven [23,73]. As well, misleading trade-off patterns may arise if more basally tolerant organisms require more extreme temperatures/longer acclimation to induce plasticity (a “threshold shift” as described in: [23]). Further, some metrics of thermotolerance— like survivorship and CCRT— are bounded, such that there is an upper and/or lower limit to said metric (e.g. 100% survival or a CCRT of 0 s). When testing for a trade-off using bounded metrics, trade-off patterns may arise due to organisms with high basal tolerance hitting the upper and/or lower limit of the metric after acclimation, rather than due to a trade-off [23,74]. The above issues, as well as others, are discussed in detail by van Heerwaarden and Kellermann in their recent review of heat tolerance-plasticity trade-off studies [23]. Considering the multitude of issues with current approaches at testing for a thermotolerance-plasticity trade-off, a reconsideration of methodology– statistical and otherwise– is vital if there is to be any hope of determining whether such a trade-off truly exists.

### Conclusions

Here, we found a strong positive correlation between basal cold tolerance and plasticity, demonstrating apparent support for a cold tolerance-plasticity trade-off at the individual level in *Drosophila melanogaster*. However, repeatability of CCO and CCRT in non-cold-acclimated flies was low, indicating that estimates of basal tolerance were unreliable, and we should therefore not be able to detect such a relationship. We argue that this pattern is likely a manifestation of regression to the mean and is not reflective of a true trade-off. Concerningly, many previous thermotolerance-plasticity trade-off studies, despite employing similar methodology to that carried out herein— regressing plasticity on basal tolerance, where plasticity is measured as the difference between basal and induced tolerance—, do not consider either repeatability or RTM in their experimental design/analyses. Moving forward, we recommend two courses of action: first, we must determine whether previous thermotolerance-plasticity trade-off studies are indeed biased by RTM (and if, in fact, there is no or little existing evidence for this hypothesis); second, we must carry out future thermotolerance-plasticity trade-off studies with appropriate controls such that this bias is accounted for in the future.

## Acronyms

CCO: chill-coma onset
CCRT: chill-coma recovery time
RTM: regression to the mean
CT_min_: critical thermal minimum
LLT: lower lethal temperature
LTL: lower thermal limit
LMM: linear mixed-effects model

## Acknowledgements

The authors wish to thank the other MacMillan lab members for their unfailing support.

## Author Contributions

All authors together conceived the study and designed the experiments. E.O. carried out the experiments. E.O. ran the systematic literature search and developed Tables 1 and 2. E.O. curated and analyzed the data and created visualizations. H.A.M. provided resources and supervision. E.O. drafted the manuscript and all authors edited the manuscript.

