## Supplementary Material for "A lack of repeatability creates the illusion of a trade-off between basal and plastic cold tolerance"

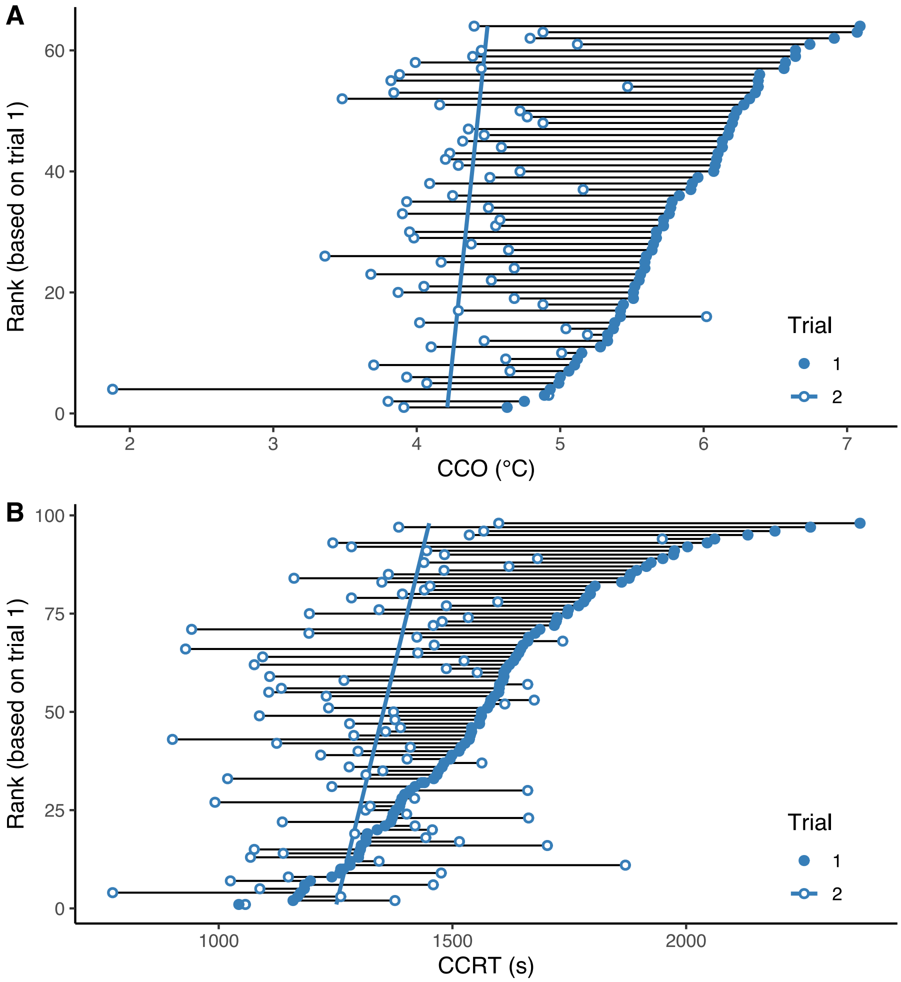


**Figure S1. 15°C-acclimated female *D. melanogaster* with greater CCO (A) or CCRT (B) in trial 1 tended to have a greater "acclimation capacity" ($CCO_1$–$CCO_2$ or $CCRT_1$–$CCRT_2$)) than individuals with lower trial 1 measurements**. Horizontal black lines connect an individual's trial 1 measurement ($CCO_1$ or $CCRT_1$; closed circles) to the same individual's trial 2 measurement ($CCO_2$ or $CCRT_2$; open circles). A lower rank on the y axis indicates higher trial 1 (basal) thermotolerance. The solid blue line is fitted to the trial 2 measurements (open circles) such that horizontal (as opposed to vertical) deviations are minimized.

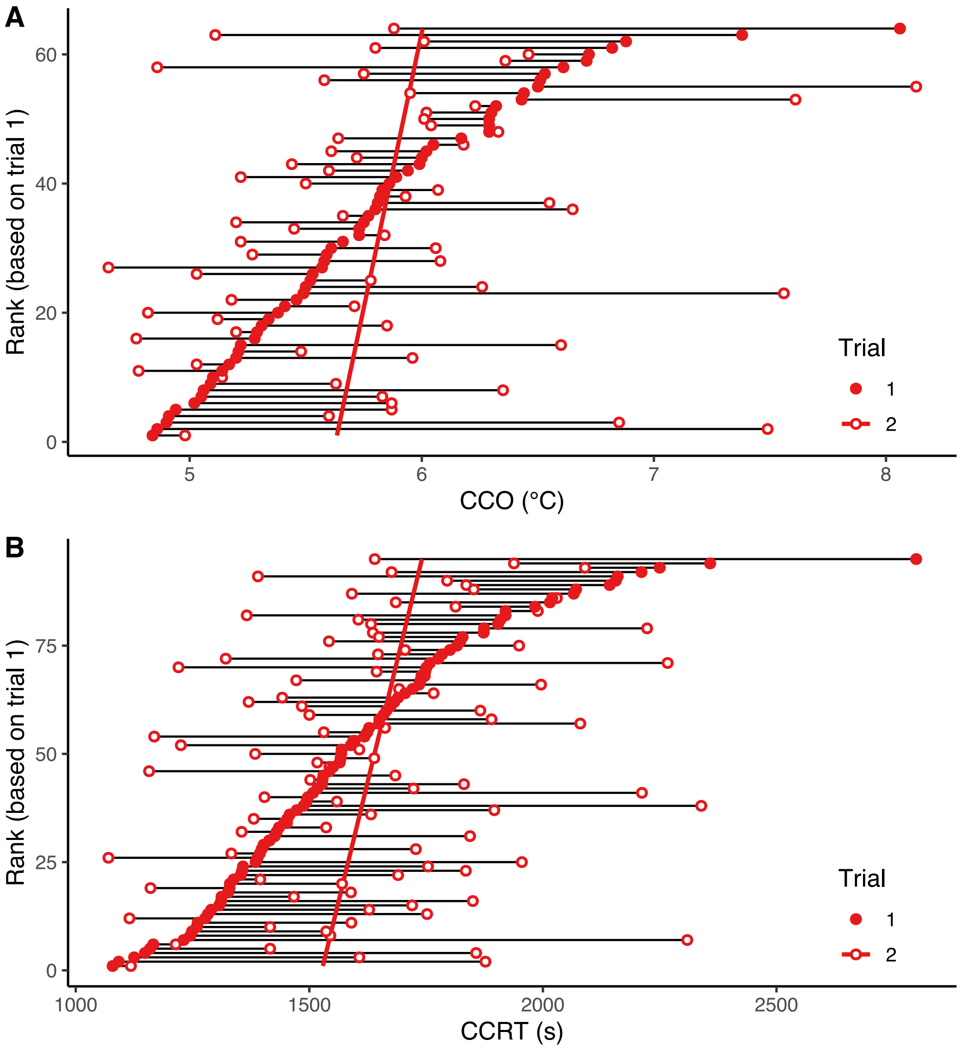


**Figure S2**. **25°C-acclimated female *D. melanogaster* that were more thermotolerant (lower CCO (A) or CCRT (B)) in trial 1 tended to be less thermotolerant in trial 2, and vice versa**. Horizontal black lines connect an individual's trial 1 measurement ($CCO_1$ or $CCRT_1$; closed circles) to the same individual's trial 2 measurement ($CCO_2$ or $CCRT_2$; open circles). A lower rank on the y axis indicates higher trial 1 thermotolerance. The solid red line is fitted to the trial 2 measurements (open circles) such that horizontal (as opposed to vertical) deviations are minimized.

**Search protocol for a review of the within-individual repeatability of thermotolerance metrics in ectotherms (see Table 2):**

**Table S1.** Search terms for a review of the within-individual repeatability of thermotolerance metrics in ectotherms.

| Search component | Search string |
|---|---|
| **Repeatability** | (repeatabilit* OR repeatable OR intraindividual OR intra-individual OR within-individual) |
| | AND |
| **Thermotolerance metric** | (thermotoleran* OR thermal-toleran* OR temperature-toleran* OR cold-toleran* OR chill-toleran* OR cold-stres* OR chill-stres* OR cold-injur* OR chill-injur* OR cold-shock OR heat-toleran* OR heat-stres* OR heat-injur* OR heat-shock OR CTmax OR CTmin OR critical-thermal OR CCO OR chill-coma OR CCRT OR SCP OR supercooling-point* OR super-cooling-point* OR knock-down-temperature* OR knockdown-temperature* OR knock-down-time* OR knockdown-time* OR loss-of-equilibrium OR LOE) |

The following databases were included in the literature search:

1. PubMed

| Search string | Details | Returns [Date] |
|---|---|---|
| (repeatabilit*[Text Word] OR repeatable[Text Word] OR intraindividual[Text Word] OR intra-individual[Text Word] OR within-individual[Text Word]) AND (thermotoleran*[Text Word] OR thermotolerance[Mesh] OR thermal-toleran*[Text Word] OR temperature-toleran*[Text Word] OR cold-toleran*[Text Word] OR chill-toleran*[Text Word] OR cold-stres*[Text Word] OR chill-stres*[Text Word] OR cold-injur*[Text Word] OR chill-injur*[Text Word] OR cold-shock[Text Word] OR heat-toleran*[Text Word] OR heat-stres*[Text Word] OR heat-injur*[Text Word] OR heat-shock[Text Word] OR CTmax[Text Word] OR CTmin[Text Word] OR critical-thermal[Text Word] OR CCO[Text Word] OR chill-coma[Text Word] OR CCRT[Text Word] OR SCP[Text Word] OR supercooling-point*[Text Word] OR super-cooling-point*[Text Word] OR knock-down-temperature*[Text Word] OR knockdown-temperature*[Text Word] OR knock-down-time*[Text Word] OR knockdown-time*[Text Word] OR loss-of-equilibrium[Text Word] OR LOE[Text Word]) | • All years (1984-2020)<br>• Advanced search<br>• *Text word and MeSH fields*<br>• Filter(s): Other animals<br>• All article types<br>• All languages<br>• English-only search terms | 50 [July 9, 2020] |

2. Web of Science

| Search string | Details | Returns [Date] |
|---|---|---|
| TS=((repeatabilit* OR repeatable OR intraindividual OR intra-individual OR within-individual) AND (thermotoleran* OR thermal-toleran* OR temperature-toleran* OR cold-toleran* OR chill-toleran* OR cold-stres* OR chill-stres* OR cold-injur* OR chill-injur* OR cold-shock OR heat-toleran* OR heat-stres* OR heat-injur* OR heat-shock OR CTmax OR CTmin OR critical-thermal OR CCO OR chill-coma OR CCRT OR SCP OR supercooling-point* OR super-cooling-point* OR knock-down-temperature* OR knockdown-temperature* OR knock-down-time* OR knockdown-time* OR loss-of-equilibrium OR LOE)) | • All years (1900-2020)<br>• Web of Science Core Collection<br>• Advanced search<br>• *Topic field*<br>• All languages<br>• All document types<br>• English-only search terms<br>• Institution subscriptions:<br>o Science Citation Index Expanded (1900-present)<br>o Social Sciences Citation Index (1956-present)<br>o Arts & Humanities Citation Index (1975-present)<br>o Conference Proceedings Citation Index - Science (1990-present)<br>o Conference Proceedings Citation Index - Social Science & Humanities (1990-present)<br>o Book Citation Index - Science & Social Science (2008-present)<br>o Current Chemical Reactions (2008-present)<br>o Index Chemicus (2008-present) | 226 [July 9, 2020] |

The structure of this search protocol is based loosely on the protocol outlined in Rytwinski et al. (2017) and the supplementary material for the accompanying review (Algera et al., 2020).
